# Protein superfolds are characterised as frustration-free topologies: A case study of pure parallel *β*-sheet topologies

**DOI:** 10.1101/2024.01.05.574326

**Authors:** Hiroto Murata, Kazuma Toko, George Chikenji

## Abstract

A protein superfold is a type of protein fold that is observed in at least three distinct, non-homologous protein families. Structural classification studies have revealed a limited number of prevalent superfolds alongside several infrequent occurring folds, and in *α/β* type superfolds, the C-terminal *β*-strand tends to favor the edge of the *β*-sheet, while the N-terminal *β*-strand is often found in the middle. The reasons behind these observations, whether they are due to evolutionary sampling bias or physical interactions, remain unclear. This article offers a physics-based explanation for these observations, specifically for pure parallel *β*-sheet topologies. Our investigation is grounded in three established structural rules that are based on physical interactions. We have identified “frustration-free topologies” which are topologies that can satisfy all three rules simultaneously. In contrast, topologies that cannot are termed “frustrated topologies.” Our findings reveal that frustration-free topologies represent only a fraction of all theoretically possible patterns, these topologies strongly favor positioning the C-terminal *β*-strand at the edge of the *β*-sheet and the N-terminal *β*-strand in the middle, and there is significant overlap between frustration-free topologies and superfolds. We also used a lattice protein model to thoroughly investigate sequence-structure relationships. Our results show that frustration-free structures are highly designable, while frustrated structures are poorly designable. These findings suggest that superfolds are highly designable due to their lack of frustration, and the preference for positioning C-terminal *β*-strands at the edge of the *β*-sheet is a direct result of frustration-free topologies. These insights not only enhance our understanding of sequence-structure relationships but also have significant implications for de novo protein design.

**Author summary:** A protein superfold is a protein fold that appears in at least three different non-homologous protein families. Superfolds are unique in their ability to accommodate multiple functions within a single fold, a feature not typically seen in other folds. Studies in structural classification have led to two notable observations: the existence of a limited number of common superfolds contrasted with a larger variety of less frequent folds, and a recurring pattern in *α/β* type superfolds where the C-terminal *β*-strand often occupies the edge of the *β*-sheet, while the N-terminal *β*-strand is usually found in the middle. The origins of these patterns, whether they stem from evolutionary sampling bias or physical interaction mechanisms, remain unclear. This article provides a physics-oriented explanation for these observations, specifically concentrating on pure parallel *β*-sheet topologies. The insights gained from this research are crucial in enhancing our understanding of the relationship between protein sequences and structures, and are expected to contribute significantly to the de novo design of new proteins.

## 1 Introduction

Research in protein structural classification has significantly advanced our understanding of the relationships between sequence, structure, function, and evolution [1–5]. These studies have also raised important new questions. One such question pertains to the relationship between protein folds and superfamilies. A protein fold is defined by the order and orientation of secondary structure elements, while a superfamily represents the largest grouping of proteins for which common ancestry can typically be inferred based on structural and functional similarity [1]. Orengo *et al*. reported that many protein folds are exclusive to a single superfamily [6], the majority of protein folds are unique to a specific superfamily. Conversely, a very small number of distinct protein folds were found across multiple superfamilies. These distinctive protein folds, observed in three or more superfamilies, are termed superfolds [6]. Interestingly, despite the existence of only nine superfolds, as many as 30% of the domains cataloged in CATH are part of these superfolds. This observation was first documented in 1994, and this trend has continued through 2021 [4]. The reasons behind the popularity of these superfolds remain an enigma.

Several structural characteristics of superfolds have been proposed to date, including pronounced symmetry [7], a significant presence of super secondary structure [8], and a limited number of jumps in *β*-sheet proteins [9]. However, it has been observed that there are folds that meet all these criteria and are thus considered potential superfolds, but these folds are either not prevalent or do not exist in the database [10, 11]. One such category is the reverse fold of superfolds, which is obtained by reversing the N- to C-terminal chain direction of a given fold. Past studies have reported that the reverse fold of the superfold is either non-existent in the database or exists in small numbers [12]. Jane Richardson, in her well-known 1981 review, stated, *“There must be some strong reason why it is so rare”* [13]. However, the reason remains unknown to this day. This is another unresolved issue raised by structural classification studies. Closely related to the issue of chain reversal is an interesting observation of the *αβ* type fold: the C-terminal *β*-strand is often located at the edge of the *β*-sheet, while the N-terminal *β*-strand tends to be in the middle of the *β*-sheet in the protein structure database [12]. The reason why the C-terminal *β*-strand strongly prefers the edge of the *β*-sheet is also, as far as we know, still unresolved.

The aim of this paper is to offer a physics-based perspective to address these questions. We focus exclusively on parallel *β*-sheet topologies with 3-6 *β*-strands for our case study. Initially, we utilized a recent database to conduct a statistical analysis, confirming that (1) superfolds make up only a fraction of all theoretically possible patterns, (2) superfolds tend to position the C-terminal *β*-strand at the edge of *β*-sheets, and (3) reverse superfolds are rare or non-existent. Subsequently, we propose a simple theory to elucidate the differences in physical properties between superfolds and their reverse counterparts. This theory shows that superfolds can simultaneously satisfy multiple physical rules, while reverse superfolds cannot. In this paper, we categorize folds that can simultaneously satisfy the physical rules as frustration-free folds, and those that cannot as frustrated folds. The theory further illustrates that frustration-free folds represent only a fraction of all theoretically possible patterns and explains why the C-terminal *β*-strand has a strong preference for the edge of *β*-sheets in frustration-free topologies. Lastly, we conducted a comprehensive exploration of sequences and structures using a lattice protein model, demonstrating that frustration leads to low designability, while non-frustration results in high designability. These findings suggest that superfolds are those with high designability due to the absence of frustration.

## 2 Results

### 2.1 The database analysis

This research evaluated all theoretically possible parallel *β*-sheet topologies comprising 3 to 6 *β*-strands. We computed the Occurrence Frequency of Homologous-group in a Topology (OFHT) for each topology within a protein structure database. This analysis aimed to confirm high skewness in the distribution of protein folds and the infrequency of reverse folds of superfolds. Here, OFHT is defined as the number of Homology-groups in a given topology (see the Materials and Methods section for details). This quantity is an indicator of how many diverse sequences have a given topology as their native structure. We employed the recent version of a semi-manually curated database, ECOD (version 20210511: develop280), which hierarchically classifies protein domains according to homology, reflecting their evolutionary relationship [14]. Utilizing the STRIDE program [15], we identified secondary structures and hydrogen bonds in the protein domains in this dataset. Following previous studies [9, 10, 12, 16], we defined *β*-sheet topologies in an abstract manner based on their *β*-strand connectivity and hydrogen bonding pattern, i.e., the number, order, and orientation of constituent *β*-strands in a *β*-sheet. These *β*-strands are sequentially numbered along the protein’s backbone, starting from the N-terminus. A topology is characterized by the *β*-strand’s sequential positions and directional orientation. Assuming that all *βαβ*-units are right-handed, there are *n*!*/*2 theoretically possible pure parallel *β*-sheet topologies consisting of *n β*-strands, so the total number of 3-6 parallel *β*-sheet topologies is 435. Out of these, 167 topologies exhibit no clash between crossing connections.

This study involves an analysis of the OFHTs in 167 clash-free topologies. The analysis reveals a highly skewed distribution, aligning with previous research [6]. In Fig. 1A, the vertical axis represents the number of topologies with a given OFHT (indicated by the horizontal axis). The distribution underscores a limited number of frequently occurring topologies alongside a larger quantity of less common ones. Following the terminology of Ref. [6], topologies with three or more OFHTs are termed “superfolds.” Six superfolds are identified in descending order of OFHT values: 2*_↑_*1*_↑_*3*_↑_*4*_↑_*, 3*_↑_*2*_↑_*1*_↑_*4*_↑_*5*_↑_*, 2*_↑_*1*_↑_*3*_↑_*4*_↑_*5*_↑_*, 3*_↑_*2*_↑_*1*_↑_*4*_↑_*5*_↑_*6*_↑_*, 1*_↑_*2*_↑_*3*_↑_* and 2*_↑_*1*_↑_*3*_↑_*, as depicted in the inset of Fig. 1A. The dataset comprises 36 normal folds (defined as topologies with OFHT values between 0 and 3) and 125 unobserved folds. It is noteworthy that these findings are based solely on pure parallel *β*-sheet proteins but are consistent with previous observations encompassing all classes of protein folds [6]. We examined whether the previous assertion regarding the rarity of reverse folds of superfolds remains valid within the current database. Our analysis of six superfolds in this dataset has revealed five topologies that transform into a different topology when the direction of the N- to C-terminal chain is reversed. The only exception is 1*_↑_*2*_↑_*3*_↑_*, which retains its topology when reversed; hence, we exclude it from further consideration. We present a comparison of the OFHTs for these five pairs of superfolds and their corresponding reversals in Fig. 1B–F. In all of these pairs, reverse folds occur infrequently; they are either entirely absent or, at most, represent less than 1/7 of the corresponding superfold, aligning with prior findings [12]. Furthermore, as depicted in Fig. 1B–F, the C-terminal *β*-strand of the superfold consistently resides at the edge of the *β*-sheet, while the N-terminal *β*-strand tends to occupy the middle of the *β*-sheet, in line with previous research [12]. We calculated the probabilities of the N- or C-terminal *β*-strand being positioned at the edge of the *β*-sheet for all observed topologies in this dataset, yielding percentages of 15.1% for the N-terminal *β*-strand and 89.1% for the C-terminal *β*-strand (Fig. 1G). In contrast, assuming that 167 clash-free topologies occur with equal likelihood, both the N- and C-terminal *β*-strands are located at the end of the *β*-sheet with a probability of 37.1%. These values differ significantly from the results of the database analysis, suggesting a compelling rationale behind the strong preference of the C-terminal *β*-strand for the edge of the *β*-sheet.

**Fig 1.**
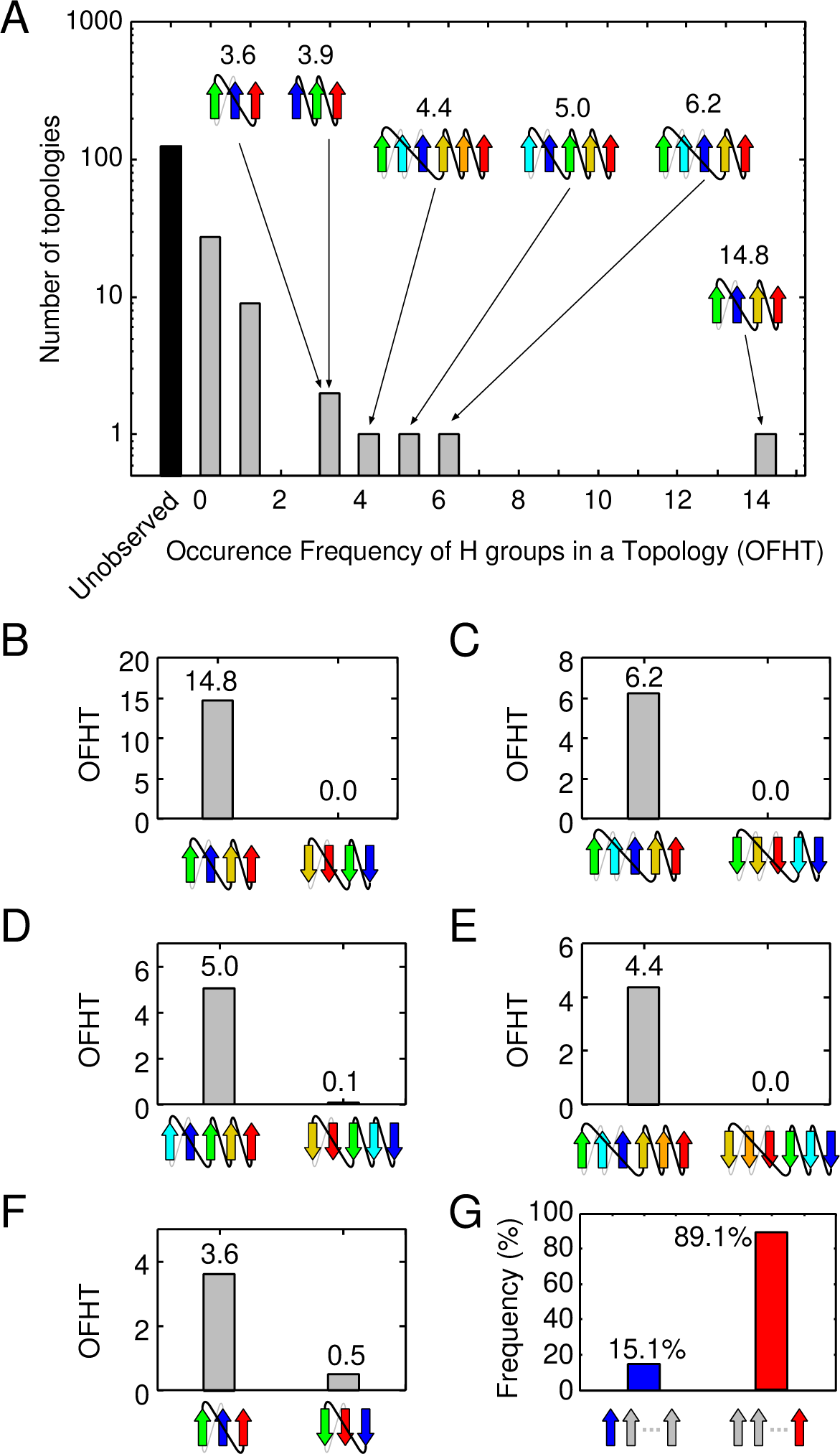
Occurrence frequency of homologous-group in a topology (OFHT) for each topology in the ECOD database. (A) It presents the distribution of pure parallel *β*-sheet topologies. The inset shows the topology diagrams of superfolds. Gray and black bars represent observed and unobserved topologies, respectively. (B)-(F) These panels depict the occurrence frequency of a superfold (on the left) and its reverse topology (on the right). (G) This panel calculates the probabilities of the N- and C-terminal *β*-strand being positioned at the edge of the *β*-sheet for all observed topologies.

### 2.2 The theory for explaining the differences in physical properties between superfolds and their reverse folds

This section introduces a theory that provides a physical explanation for why a superfold is more favorable than its reverse fold. The term “physically favorable” refers to a topology’s ability to simultaneously comply with three specific rules outlined below.

#### Rule I. The right-handed rule for crossover connections of βαβ-unit

Crossover connections in the *βαβ*-unit predominantly exhibit right-handedness [17, 18]. This rule applies to the two connected *β*-strands that are nearest neighbors in the *β*-sheet and to the two connected *β*-strands that have one or more intervening *β*-strands [17]. The rationale for this rule lies in thermodynamic stability and kinetic accessibility [19–21]. Notably, over 98% of the *βαβ*-motifs in the database display right-handedness [21], a finding corroborated by our recent calculations (Fig. 2A). For detailed methods on calculating the handedness of *βαβ*-units, please refer to Ref. [22].

**Fig 2.**
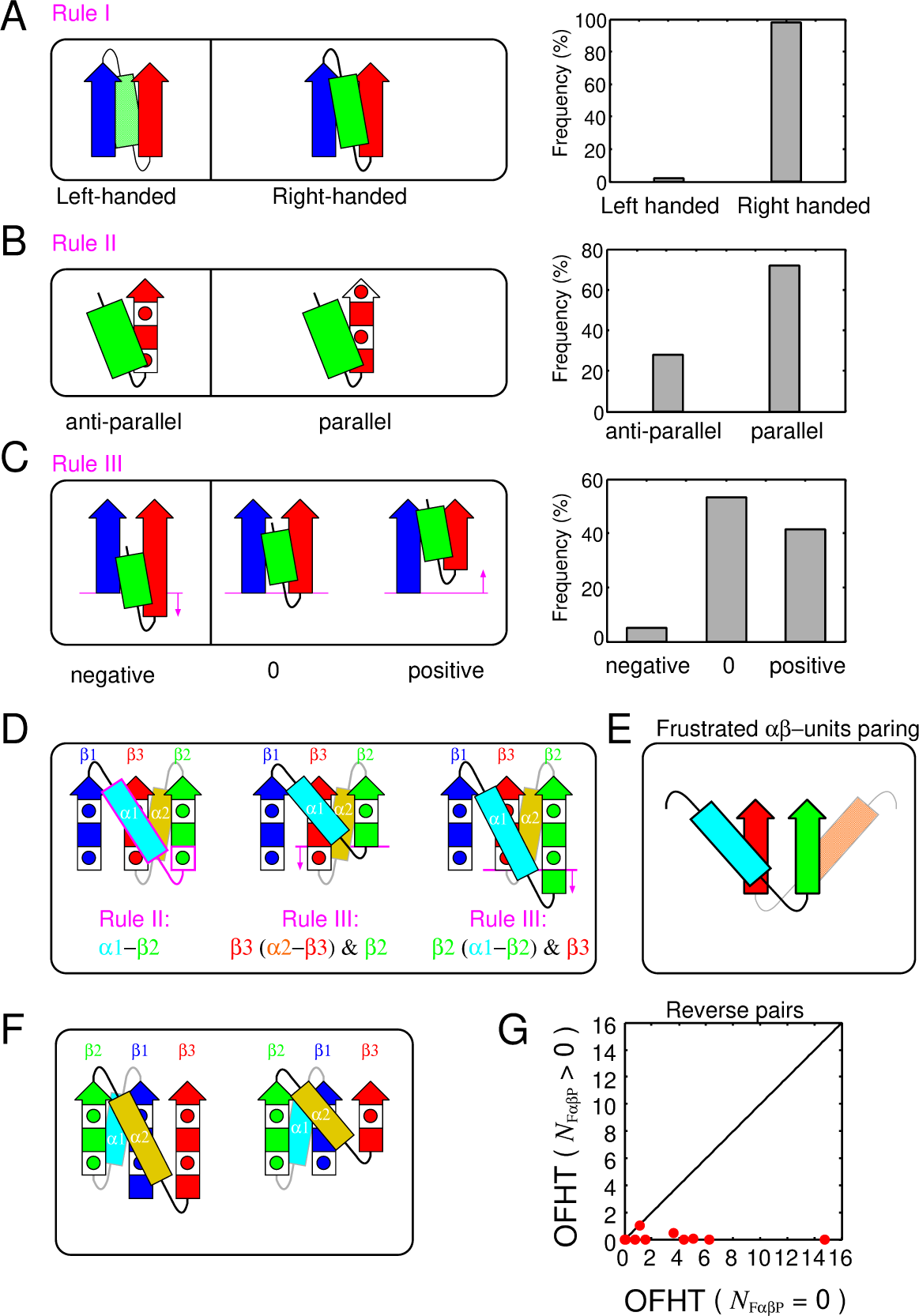
The basic rules and the frustrated *αβ*-units paring. (A) The right-handed rule for crossover connections of *βαβ*-unit is depicted. The left side shows schematics of left- and right-handed *βαβ*-units, while the right side presents their occurrence frequencies. (B) The *αβ*-rule is represented. The left side displays schematics of antiparallel and parallel *αβ*-units, and the right side shows their occurrence frequencies. The square with a circle inside symbolizes a single amino acid residue with a side chain located on the proximal side, and the red-colored filled square represents a residue with a side chain located on the opposite side. (C) The register shift rules for a *β*-strand of an *αβ*-unit and its neighboring *β*-strand are depicted. The left side provides a schematic representation of negative, zero, and positive register shifts, and the right side shows their occurrence frequencies. (D) A topology that cannot simultaneously satisfy the three rules is illustrated. (E) A schematic of frustrated *αβ*-units paring is shown. (F) A topology that can simultaneously satisfy the three rules is depicted. (G) OFHTs of topology pairs are presented, where one is a topology that does not contain a F*αβ*P and the other is its reverse topology that does.

#### Rule II. The αβ-rule

The *αβ*-units generally exhibit a parallel orientation [23]. This means that the vector from the *α*-helix to the *β*-strand of an *αβ*-unit aligns parallel to the C*α*-C*β* vector of the first residue in the strand. When the two are antiparallel, the orientation of *αβ*-units becomes antiparallel (see Fig. 2B). This rule, discovered through database analysis and confirmed in physics-based simulations, is believed to result from physical interactions [23]. This rule is not as strict as the right-handed rule of *βαβ*-units, with a ratio of approximately 7:3 for parallel to antiparallel orientations (see Fig. 2B).

#### Rule III. The register shift rules for a β-strand of an αβ-unit and its neighboring β-strand

The register shift between the N-terminal residue of the *β*-strand of the *αβ*-unit and the N-terminal residue of its neighboring *β*-strand must not be negative [24]. To clarify, the residue offset between the N-terminal residue of the *β*-strand of the *αβ*-unit and the N-terminal residue of the *β*-strand to its left when viewed from a specific direction, should not be negative [24]. This specific direction is defined as the one where the *α*-helix of the *αβ*-unit is closer to the *β*-strand and the *β*-strand is facing upwards (Fig. 2C). The sign of the register shift is defined as positive when the N-terminal residue of the *β*-strand of the *αβ*-unit is located above the N-terminal residue of the *β*-strand to its left, and negative when the opposite is the case (see the left side of Fig. 2C). For a detailed and comprehensive definition, please refer to Ref. [24]. In this context, “negative register shifts” are strongly discouraged, as indicated by database analysis (see the right side of Fig. 2C). The physical mechanism that prevents negative register shifts has been clarified through all-atom model calculations and exhaustive structure sampling [24]. It is important to note that the blue *β*-strand and the green *α*-helix shown in Fig. 2C do not need to be directly connected [24].

In this section, we clarify how certain topologies can simultaneously satisfy the three aforementioned rules, while others cannot. We use 2*_↑_*1*_↑_*3*_↑_*, depicted on the left side of Fig. 1F, as an example of a topology that can comply with all three rules, and 1*_↑_*3*_↑_*2*_↑_* (equivalent to 2*_↓_*3*_↓_*1*_↓_*), also shown on the right side of Fig. 1F, as an example of a topology that cannot. It is important to note that these two topologies are reverses of each other. In the ensuing discussion, we will assume that the right-handed rule of the *βαβ*-motif always holds. We will refer to the *β*-strands in these topologies as *β*1, *β*2, and *β*3, starting from the N-terminal side. The *α*-helix connecting *β*1 and *β*2 will be referred to as *α*1, and the one connecting *β*2 and *β*3 as *α*2 (see Fig. 2D and F). Firstly, we will explain why 1*_↑_*3*_↑_*2*_↑_*cannot simultaneously adhere to all three rules in three steps.

##### 1st step

When the N-terminal residues of *β*2 and *β*3 are positioned at the same height as shown on the left side of Fig. 2D, neither the *αβ*-unit consisting of *α*1 and *β*2 nor the one consisting of *α*2 and *β*3 can satisfy Rule II. As illustrated on the left side of Fig. 2D, the *αβ*-unit consisting of *α*1 (cyan) and *β*2 (green) violates Rule II, as the vector from the *α*-helix to the *β*-strand is antiparallel to the C*α*-C*β* vector of the first residue of *β*2 (the region breaking the rule is outlined in magenta). Conversely, if the N-terminal side chains of *β*2 and *β*3 are on the other side of the paper, the *αβ*-units consisting of *α*2 (yellow) and *β*3 (red) violate Rule II. Therefore, when the register shift of the N-terminal side of *β*2 and *β*3 is zero, Rule II cannot be satisfied, regardless of the orientation of the side chains of the N-terminal residues of *β*2 and *β*3.

##### 2nd step

If the N-terminal residue of *β*2 is located above the N-terminal residue of *β*3, Rule III cannot be satisfied because the register shift between *β*2 and the *β*-strand of the *αβ*-unit consisting of *α*2 and *β*3 is negative (see the middle of Fig. 2D).

##### 3rd step

If the N-terminal residue of *β*2 is located below the N-terminal residue of *β*3, Rule III cannot be satisfied because the register shift between *β*3 and the *β*-strand of the *αβ*-unit consisting of *α*1 and *β*2 is negative (see the right side of Fig. 2D).

Considering these three steps, the 1*_↑_*3*_↑_*2*_↑_*topology cannot simultaneously satisfy all three rules for any offset value between the N-terminal residues of *β*2 and *β*3. Broadening the scope of the discussion, a topology cannot meet all three rules if two *αβ*-units form parallel *β*-sheets, and each *α*-helix is on a different *β*-sheet face side, as depicted in Fig. 2E. We refer to such a pair of *αβ*-units as a frustrated *αβ*-units pairing (F*αβ*P) and denote the number of F*αβ*P in a topology as *N*_F*αβ*P_.

Next, we explain how the 2*_↑_*1*_↑_*3*_↑_* topology can simultaneously fulfill all three rules. This topology does not involve F*αβ*P, indicating that it can satisfy all three rules simultaneously. In fact, there are certain arrangements of *β*-strands that can meet all three rules. Fig. 2F provides two examples of such *β*-strand arrangements for the 2*_↑_*1*_↑_*3*_↑_* topology.

The observation that a superfold lacks F*αβ*P, while its reversal with infrequent occurrences contains it, holds true for all pairs of superfolds and their reversals, not just the specific pair of 1*_↑_*3*_↑_*2*_↑_*and 2*_↑_*1*_↑_*3*_↑_* topologies. It can be readily confirmed that the superfolds presented in Fig. 1B–F also lack F*αβ*P, while their reversals contain it. This observation suggests that the absence of F*αβ*P is one of the necessary conditions for a topology to be classified as a superfold. This implication is further supported by the distribution data obtained from topologies with *N*_F*αβ*P_ = 0 and those with *N*_F*αβ*P_ *>* 0; all superfolds are found within the category of topologies with *N*_F*αβ*P_ = 0, whereas topologies with *N*_F*αβ*P_ *>* 0 are not designated as superfolds (S1 FigA and B). However, it is important to emphasize that having *N*_F*αβ*P_ = 0 alone does not suffice as a condition for a topology to be recognized as a superfold.

The condition *N*_F*αβ*P_ = 0 is not the only criterion for classifying a topology as a superfold; there are other factors to consider. One such factor is the small number of jumps (*N*_j_) (S1 FigC and D). *N*_j_ is defined as the count of pairs of *β*-strands that are adjacent in sequence but not neighboring in the *β*-sheet structure. Prior studies have emphasized that topologies with more than one jump (*N*_j_ *>* 1) are strongly discouraged within the Protein Data Bank (PDB) [9]. This discouragement stems from the fact that topologies with a high number of jumps encounter significant challenges during the folding process, primarily due to the substantial free energy barrier resulting from rapid entropy loss [25]. Therefore, both the number of frustrated *αβ*-units pairings and the number of jumps are key determinants in the classification of a topology as a superfold.

However, it is important to emphasize that only the former factor, the number of F*αβ*Ps, can provide insights into the differences in physical properties between pairs with reversed chain orientations, potentially influencing the OFHTs, as illustrated in Fig. 1B–F. This relationship extends beyond these reversed pairs and applies to all such pairs. Among all theoretically possible topologies, there are 23 topology pairs where one lacks F*αβ*P, while its reverse contains F*αβ*P. Fig. 2G presents a graphical representation of the OFHTs for these 23 topology pairs. In this plot, each pair’s OFHT with *N*_F*αβ*P_ = 0 is plotted on the horizontal axis, while the OFHT of its reverse topology with *N*_F*αβ*P_ *>* 0 is depicted on the vertical axis. Notably, all data points on the plot are located exclusively on the lower right-hand side of the diagonal line. This observation underscores the significant impact of the presence of F*αβ*P on the reduction of OFHTs.

### 2.3 Superfolds are characterised as frustration-free topologies

In this section, we undertake a more comprehensive characterization of superfolds, focusing on two pivotal factors: the number of F*αβ*P and the number of jumps within a *β*-sheet. Fig. 3A–D depict the distribution of topologies across four distinct categories: theoretically possible crash-free topologies, superfolds, normal folds, and unobserved folds. These topologies are charted on a 3 *×* 6 two-dimensional grid, where the horizontal axis signifies the number of F*αβ*Ps (*N*_F*αβ*P_), and the vertical axis represents the number of jumps (*N*_j_). Notably, states highlighted in red denote instances where over 50% of the theoretically possible crash-free topologies within the states fall into the categories of superfolds (Fig. 3B), normal folds (Fig. 3C), or unobserved folds (Fig. 3D). The initial salient observation is that the number of topologies meeting both physically preferred conditions, *N*_F*αβ*P_ = 0 and *N*_j_ *≤* 1, is notably lower than the total count (Fig. 3A). This implies a constrained number of topologies adhering to physically favorable criteria. The second critical insight is that all six superfolds are concentrated within states characterized by *N*_F*αβ*P_ = 0 and *N*_j_ *≤* 1 (Fig. 3B). Conversely, topologies falling within grids defined by *N*_F*αβ*P_ = 0 and *N*_j_ *≤* 1 are likely to be superfolds, with 6 out of 14 topologies being superfolds. Remarkably, within the grid where (*N*_F*αβ*P_ = 0 and *N*_j_ = 1), as many as 50% (5 out of 10) of topologies are superfolds. These observations suggest that superfolds can be effectively characterized as topologies that satisfy all physically favorable requirements (*N*_F*αβ*P_ = 0 and *N*_j_ *≤* 1). Henceforth, we will denote a topology meeting all these criteria (*N*_F*αβ*P_ = 0 and *N*_j_ *≤* 1) as a frustration-free topology, while one that fails to meet these criteria will be termed a frustrated topology. The third crucial point is that as topologies deviate further from the state of (*N*_F*αβ*P_*, N_j_*) = (0, 1), there is a discernible decrease in the overall OFHT (Fig. 3C and D). This implies that both of these variables contribute significantly to distinguishing between superfolds, normal folds, and unobserved folds. Based on these findings, we propose the “frustration-free hypothesis of superfolds”: a superfold is a topology that impeccably satisfies all various physical rules. Subsequently in this paper, we will present data to support this hypothesis from a lattice protein model study.

**Fig 3.**
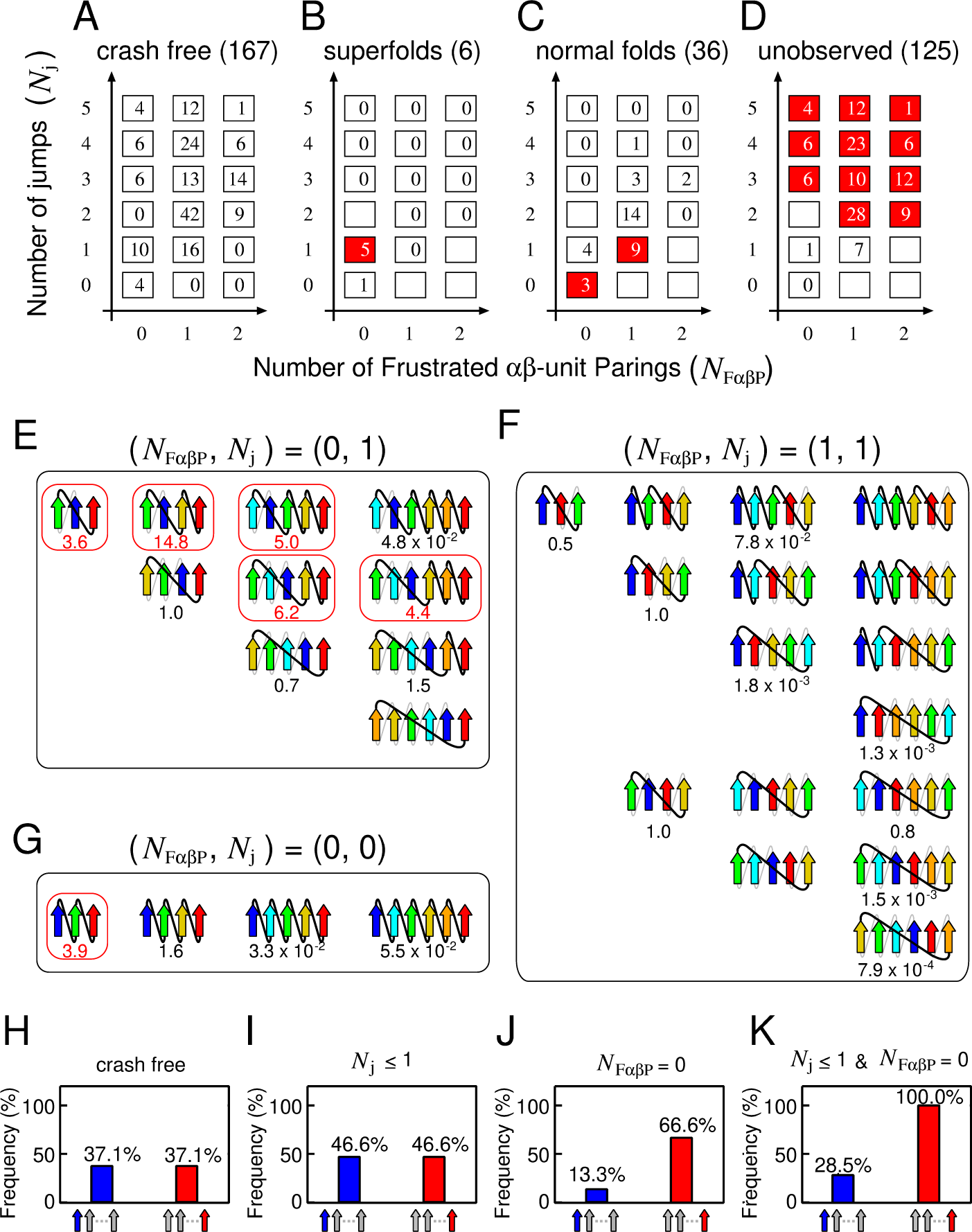
Topologies categorized by *N*_F*αβ*P_ and *N*_j_. (A)-(D): The number of topologies for theoretically possible crash-free states, superfolds, normal folds, and unobserved folds, depicted on a two-dimensional grid. The states highlighted in red indicate instances where more than 50% of the total number of theoretically possible crash-free topologies are observed. (E)-(G): Topology diagrams in the states (*N*_F*αβ*P_*, N_j_*) = (0, 1), (*N*_F*αβ*P_*, N_j_*) = (1, 1), and (*N*_F*αβ*P_*, N_j_*) = (0, 0). The numbers below the topology diagrams represent their OFHTs. Topology diagrams surrounded by a red box represent superfolds. (H)-(K): The probabilities of the N- or C-terminal *β*-strand being at the edge of the *β*-sheet calculated from crash-free topologies, topologies with (*N*_j_ *≤* 1), those with (*N*_F*αβ*P_ = 0), and those with (*N*_j_ *≤* 1, *N*_F*αβ*P_ = 0).

In this study, we delve into the characteristics of topologies within different grid states. We specifically present the topologies in three particular grids: (*N*_F*αβ*P_*, N_j_*) = (0,1), (1,1), and (0,0). However, the topologies of the other grids are detailed in S2 Fig - S7 Fig.

The first example pertains to the state with (*N*_F*αβ*P_*, N_j_*) = (0,1). This grid exhibits the highest likelihood of presenting superfolds in this 2D plane. It includes ten clash-free topologies (Fig. 3B). Out of these, five topologies display superfolds, while the others do not. At present, the reasons for these five non-superfold topologies not exhibiting superfolds remain uncertain. One possible explanation for the absence of the 5*_↑_*4*_↑_*3*_↑_*2*_↑_*1*_↑_*6*_↑_* topology in PDB is the presence of a significant number of intervening *β*-strands between *β*-strand 5 and 6, which is anticipated to be physically unfavorable, thereby disqualifying it as a superfold. A notable structural characteristic of this group is the consistent positioning of the C-terminal *β*-strand at the edge of the *β*-sheet, while the N-terminal *β*-strand consistently resides in the middle across all topologies. This structural pattern could account for the strong preference for C-terminal *β*-strands at the edge of the *β*-sheet in PDB.

The second example pertains to the state with (*N*_F*αβ*P_*, N_j_*) = (1,1), which does not contain a superfold and has the second highest probability of a normal fold (9/16). This state comprises reverse topologies of (*N*_F*αβ*P_*, N_j_*) = (0,1) (upper 4 rows) and other topologies (lower 3 rows) (Fig. 3C). A notable feature of this state is the high probability (62.5%) of the N-terminal *β*-strand being positioned at the edge of the *β*-sheet, while the C-terminal *β*-strand consistently occupies the middle position across all topologies within this state. Notably, the difference in the physical properties of topologies with (*N*_F*αβ*P_*, N_j_*) = (0,1) and those with (*N*_F*αβ*P_*, N_j_*) = (1,1) has previously gone unrecognized. However, F*αβ*P has revealed their distinct physical characteristics. Our hypothesis suggests that topologies within this state cannot achieve superfold status due to the presence of F*αβ*P.

The third example pertains to the state with (*N*_F*αβ*P_*, N_j_*) = (0,0), which exhibits the highest likelihood of a normal fold. This state encompasses four topologies (Fig. 3D). A significant structural characteristic of this state is that both the N- and C-terminal *β*-strands consistently occupy positions at the edge of the *β*-sheet and lack non-local contacts due to *N*_j_ = 0. Despite this grid state being free of frustration, only one superfold exists within this grid. One potential explanation for the three topologies not being superfolds is their lack of non-local contacts, leading to low cooperativity as anticipated [26]. This characteristic might be a factor making them less likely to be superfolds. Determining whether the reason these three topologies are not superfold is due to evolutionary sampling bias or physical necessity is a subject for future investigation.

In the final segment of our 2D plane analysis, we demonstrate that the likelihood of the N- or C-terminal *β*-strands being located at the edge of the *β*-sheet aligns closely with database observations (Fig. 1G). This alignment is particularly noticeable when the topologies meet specific conditions that are physically favorable (*N*_j_ *≤* 1 and *N*_F*αβ*P_ = 0). In the following discussion, we assume that all topologies occur with equal probability. Fig. 3H provides a visual representation of the probability of an N- or C-terminal *β*-strand being located at the edge of the *β*-sheet across all 167 crash-free topologies. It becomes apparent that the N- and C-terminal *β*-strands have an equal probability (37.1%) of occupying the end of the *β*-sheet. Fig. 3I presents the probabilities calculated for the 30 topologies that meet the *N*_j_ *≤* 1 condition. Here, the N- and C-terminal *β*-strands have an identical likelihood of being situated at the edge of the *β*-sheet. It is clear that the *N*_j_ *≤* 1 condition alone does not influence the probability of the N- and C-terminal *β*-strand being at the edge of the *β*-sheet. When we perform similar calculations for the 30 topologies that meet the *N*_F*αβ*P_ = 0 condition (Fig. 3J), we observe a difference in the probability distribution between the N- and C-terminal *β*-strands. The C-terminal *β*-strand has a higher likelihood of being located at the edge than the N-terminal *β*-strand. This finding underscores the significant impact of the presence or absence of F*αβ*P on the preferred positioning of the N- and C-terminal *β*-strands. Topologies that satisfy both conditions, *N*_F*αβ*P_ = 0 and *N*_j_ *≤* 1 (Fig. 3K), show an even higher probability of the C-terminal *β*-strand being located at the edge. This observation aligns with the findings in Fig. 1G. Collectively, these insights suggest that the prevalence of proteins with C-terminal *β*-strands at the edge of the *β*-sheet in the database can be attributed to the preference for frustration-free topology.

### 2.4 Frustration-free structures are highly designable: Implication from a lattice model study

In the previous section, we established a connection between superfolds and frustration-free topologies, demonstrating that all superfolds are frustration-free and that frustration-free topologies are often associated with superfolds. This leads us to the question: why are frustration-free protein topologies so common among various protein families? To address this, we utilize a simplified two-dimensional lattice HP model for proteins [27], which incorporates unfavorable local interactions [28], as illustrated in Fig. 2A-C. Our investigation reveals that frustration-free structures exhibit high designability. Here, the statement that a structure is highly designable means that when all theoretically possible amino acid sequences are considered, substantial number of the sequences adopt the structure as their native state [29]. Given that a superfold is defined as a protein fold observed in a large number of non-homologous families, high designability is a prerequisite for a structure to be classified as a superfold.

In the lattice HP model, proteins are represented as self-avoiding paths on a two-dimensional square lattice, featuring two types of amino acids: hydrophobic (H) and polar (P). This simplistic representation facilitates a detailed exploration of the interplay between protein sequences and structures, especially for short protein chains. In the original HP model [27], the energy, denoted as *E*_HP_, associated with a protein chain’s conformation is solely determined by the count of hydrophobic–hydrophobic (H–H) contacts, represented as *n*_HH_. This energy is expressed as *E*_HP_ = *−ɛ*_HH_ *· n*_HH_, where *ɛ*_HH_ is a positive constant. Beyond the original energy function, we introduce a sequence-independent energy penalty to emulate the effects shown in Fig. 2A-C. We define three specific local structures, as visualized in Fig. 4A-C, as inherently unfavorable structures. Each of these structures incurs an energy penalty, *E*_penalty_ = *ɛ*_penalty_ *· n*_penalty_, where *n*_penalty_ denotes the number of such unfavorable structures within a given configuration, and *ɛ*_penalty_ is a positive constant. Consequently, the overall energy, *E*, of a given protein structure and its corresponding sequence is computed as *E* = *E*_HP_ + *E*_penalty_. Considering that the structure depicted in Fig. 4D necessarily includes one of the three structures presented in Fig. 4A-C, resembling the frustration observed in the topology shown in Fig. 2D, we term this structure a ‘Frustrated Local Structure’ (FLS). An example of a structure containing an FLS is illustrated in Fig. 4E. Note that when this structure resides at the most C-terminal end, as shown in Fig. 4F, no energetic penalty is imposed due to the absence of the structures illustrated in Fig. 4A-C. Furthermore, the structure depicted in Fig. 4F, the reverse of the Fig. 4E structure, is devoid of frustration, closely resembling the pair of topologies shown in Fig 2F and Fig. 2D.

**Fig 4.**
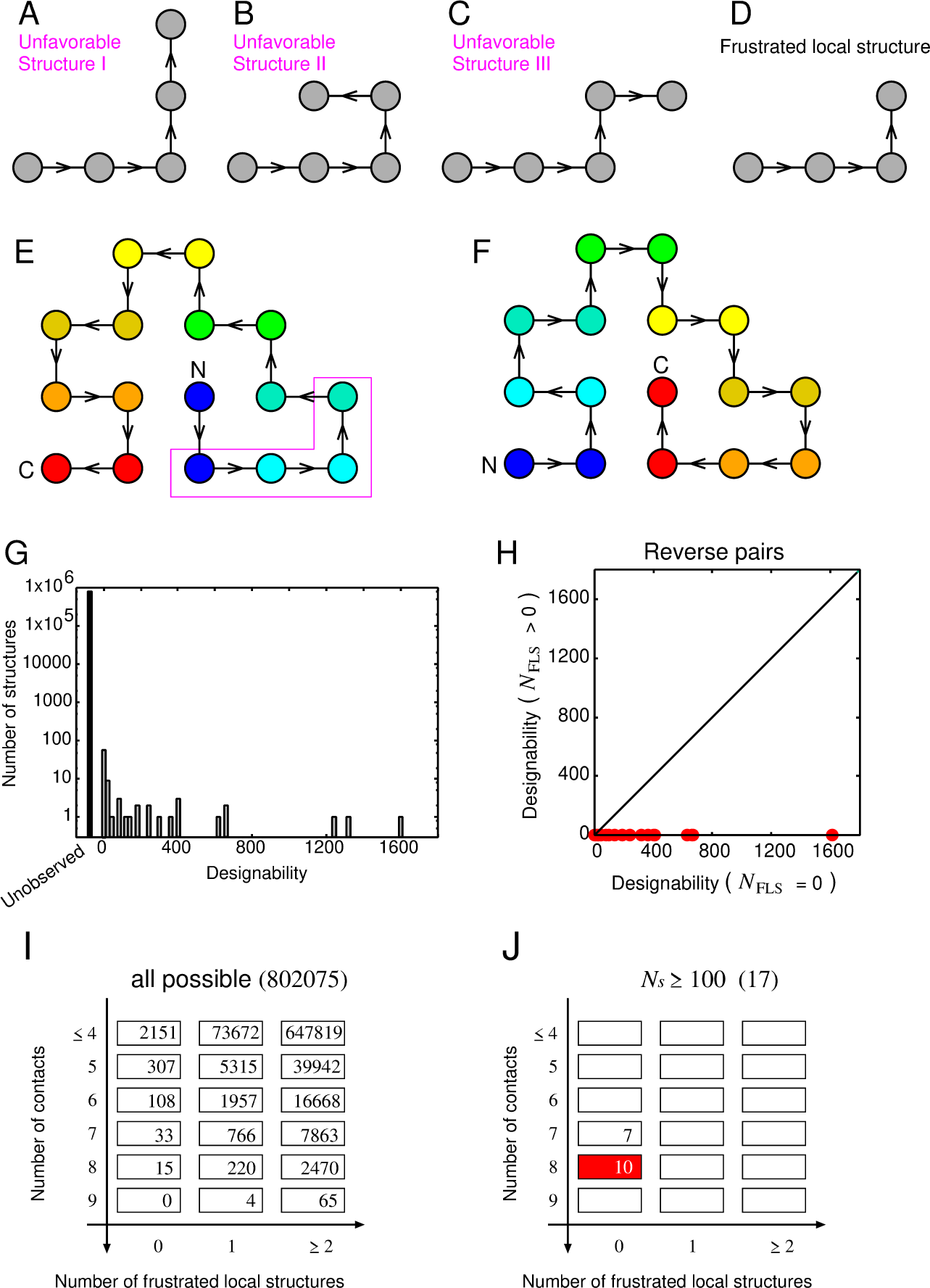
The lattice protein model employed in this study. (A)-(C) Depict three structures defined as physically unfavorable. (D) Represents frustrated local structure (FLS), a local structure that necessarily contains one of the three physically unfavorable structures. (E) Provides an example of a structure containing an FLS. Rainbow coloring from blue to red indicates the N- to C-terminal position of the residues in the model. (F) Illustrates the reverse structure of (E), which does not contain an FLS. (G) Shows a histogram of designability for the lattice model with the local interactions. (H) Displays designabilities of structure pairs, where one is a structure that does not contain an FLS and the other is its reverse structure that does. The black bar and gray bar represent the number of structures with *N_s_* = 0 and those with *N_s_* = 0, respectively. (I)-(J) Illustrates the number of all possible structures and highly designable structures mapped onto the two-dimensional grid, *N*_F*αβ*P_ and *N*_FLS_.

In this study, we conducted an analysis of protein chains composed of 16 monomers, resulting in a total of 802,075 conformations for this chain length. For each given sequence, we computed the energies of all structures using the parameters *ɛ*_HH_ = 1.0 and *ɛ*_penalty_ = 2.0. Following a common practice in other research studies, only sequences exhibiting a unique ground state are categorized as protein-like [29]. We exhaustively enumerated all possible conformations for the all possible 2^16^ sequences and identified those meeting the criteria for being protein-like, resulting in a total of 10,139 such sequences. Following this, we calculated the designability for each structure. In this context, the designability of a specific structure (*S*) is defined as the count of sequences (*N_S_*) that have the structure *S* as their unique ground state. It has been observed that in the HP model, there’s a considerable variation in the designability of structures. A small number of structures are highly designable, while a significant number of structures exhibit low designability [29].

Our model, which incorporates local interactions, mirrors the findings of numerous previous studies on lattice protein models [29–33]. It exhibits a limited number of highly designable structures and a substantial number of structures with low designability (see Fig. 4G). This observation aligns qualitatively with the data presented in Fig. 1A, suggesting that the relationship between the number of folds and families within the database can be elucidated by examining the interplay between structure and sequence in our current model. To investigate the influence of FLS on designability, we categorized all 802,075 structures into two distinct groups: those with FLS and those without FLS. We then plotted the relationship between designability and the number of structures for each group (see S8 Fig A and B). Our investigation revealed a consistent pattern: all structures characterized by high designability were devoid of FLS, while structures containing FLS consistently exhibited low designability (S8 Fig A and B). These findings suggest that the absence of FLS is a prerequisite for a structure to exhibit high designability.

While the absence of FLS is a key factor in determining high designability, it is not the only criterion. There are other contributing factors, such as the number of contacts (*N_c_*), which is another crucial determinant of high designability (S8 Fig C and D). Therefore, both presence of FLS and the number of contacts influence designability. However, the presence of FLS is the only factor that can account for differences in the designability between pairs with reversed chain orientation. For example, consider the structure with the highest designability (*N_s_* = 1614) depicted in Fig. 4F, and its reverse structure (*N_s_* = 3) shown in Fig. 4E. Both structures have the same number of contacts, making it impossible to distinguish them based solely on contact numbers. In contrast, the number of FLS can differentiate these two cases: the structure with the highest designability lacks FLS, while its reverse structure contains FLS. This relationship extends beyond this specific pair and remains consistent across all pairs (Fig. 4H). These findings underscore the pivotal role of the number of FLS as one of the most critical determinants of designability, offering unique insights compared to other established determinants.

Highly designable structures can be more effectively characterized using two specific features: the number of FLSs (*N*_FLS_) and the number of contacts (*N*_c_). Fig. 4I and 4J visually represent the number of all possible structures and highly designable structures (defined as *N*_s_ *≥* 100), mapped onto a 3 *×* 6 two-dimensional grid based on *N*_FLS_ and *N*_c_. The red-shaded region within Fig. 4J indicates the state where over 50% of all possible structures within the state are highly designable. A significant observation from Fig. 4I is that the number of structures satisfying both criteria (*N*_c_ *≥* 7 and *N*_FLS_ = 0) is substantially smaller than the total number, suggesting a limited number of physically favorable structures. Notably, all 17 highly designable structures (Fig. 4J) fall within the grids defined by (*N*_c_ *≥* 7 and *N*_FLS_ = 0). Conversely, structures located within grids meeting these criteria (*N*_c_ *≥* 7 and *N*_FLS_ = 0) are likely to be highly designable, with 17 out of 48 structures meeting this classification. The grid with (*N*_c_ = 8 and *N*_FLS_ = 0) is particularly significant, where as many as 66% (10 out of 15) structures exhibit high designability. Thus, these two variables prove to be effective in characterizing highly designable structures.

Applying the results derived from lattice model calculations to the database analysis of pure parallel *β*-sheet topologies, we can conclude that: (i) the frustration-free topologies exhibit high designability, thereby fulfilling the necessary condition for being superfolds, and (ii) the frustrated topologies cannot be designated as superfolds due to their low designability. It is crucial to underscore that the feature *N*_F*αβ*P_, which accounts for different physical properties linked to the reversal of chain direction, plays a pivotal role in addressing the two critical questions: why is the population of superfolds limited, and why do the reverse folds of the superfolds either not exist in the database at all or exist only in small numbers?

## 3 Discussions

What sets superfolds apart from other protein structures? Is their prevalence merely a result of evolutionary sampling bias, or is there a fundamental reason underlying their ubiquity? In this study, we provide compelling evidence that unequivocally sets superfolds apart as unique entities. Superfolds are characterized by their nature as frustration-free topologies, a feature that is relatively rare when compared to the vast array of all possible topologies. Notably, the identification of a frustration-free topology does not necessitate sequence information or energy calculations; it can be determined solely based on the topology and compliance with several physical rules. Our findings, derived from calculations using the HP lattice model, reveal that frustration-free structures exhibit significantly high designability. Therefore, we conclude that the widespread occurrence of superfolds across diverse protein families is primarily due to their elevated designability, rather than being merely a consequence of evolutionary sampling bias.

Among the various principles that govern protein structures, the right-handed rule for crossover connections of *βαβ*-units is perhaps the most well-known. This rule states that the majority of *βαβ*-units are predominantly right-handed [17, 18]. However, in our dataset, we have identified a small subset of left-handed *βαβ*-units, as illustrated in Fig. 2A. This raises the question: are there specific topological features where left-handed *βαβ*-units are frequently observed? To investigate this, we calculated the OFHTs for domains composed of four or more *β*-sheets containing the substructure 1*_↑_*3*_↑_*2*_↑_*or 2*_↑_*1*_↑_*3*_↑_*. For each substructure, we computed the percentage of left-handed *βαβ*-units included in those substructures (see Fig. 5A). Interestingly, the probability of encountering a left-handed *βαβ*-unit in domains containing 1*_↑_*3*_↑_*2*_↑_* as a substructure (7.5%) is approximately 100 times higher than that in domains containing 2*_↑_*1*_↑_*3*_↑_*(0.073%). This observation suggests the existence of a mechanism that favors the occurrence of left-handed *βαβ*-units in topologies containing 1*_↑_*3*_↑_*2*_↑_*. A plausible explanation for this mechanism is that domains containing 1*_↑_*3*_↑_*2*_↑_*often prioritize satisfying Rule II and Rule III over Rule I, consequently leading to the adoption of left-handed *βαβ*-units. An example of such a structure that adheres to other rules while violating Rule I is depicted in Fig. 5B. These findings suggest that in frustrated topologies, Rule I is often violated due to competition with other rules, making left-handed *βαβ*-units more likely to occur compared to frustration-free topologies.

**Fig 5.**
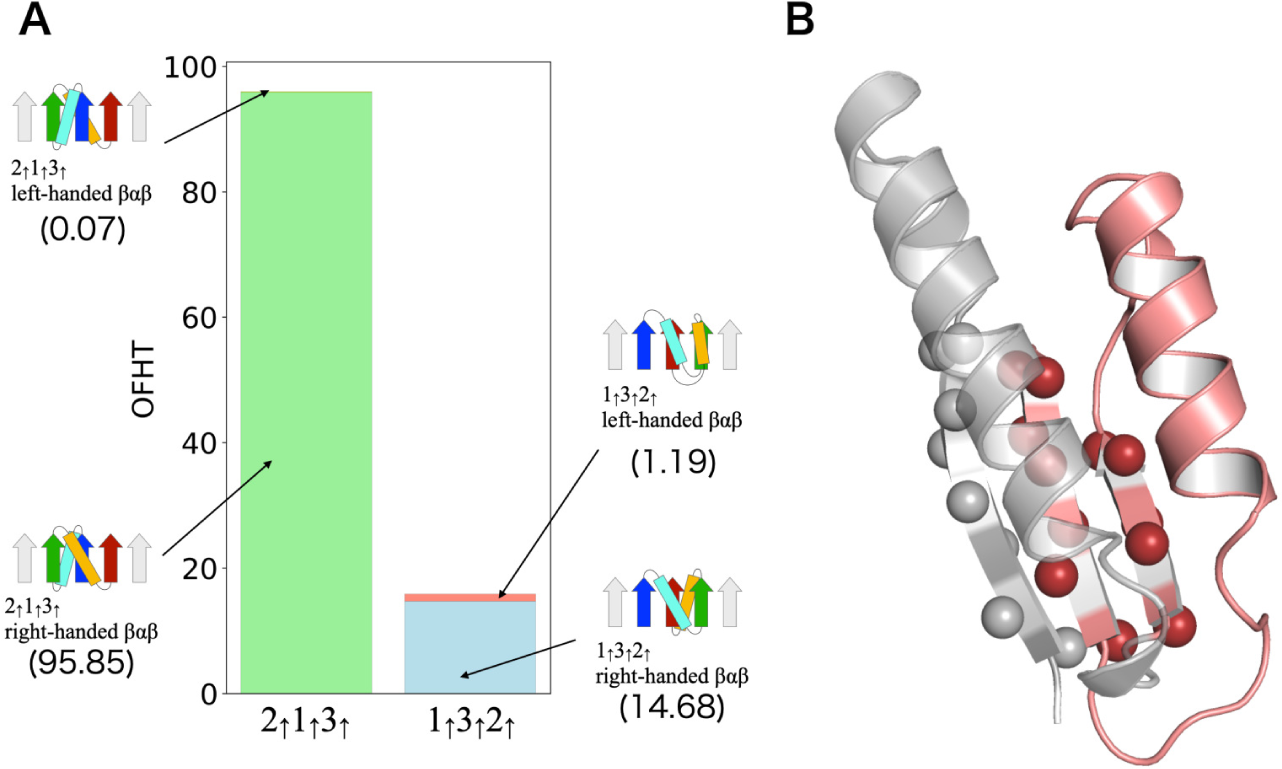
Left-handed *βαβ*-units in the dataset. (A) Depicts the occurrence frequencies of the left-handed and right-handed *βαβ*-units in the 2*_↑_*1*_↑_*3*_↑_*and 1*_↑_*3*_↑_*2*_↑_* substructures. The OFHT of topologies containing the 2*_↑_*1*_↑_*3*_↑_*substructures is 95.92 among which the OFHT of topologies having the left-handed *βαβ*-unit is 0.07. The OFHT of topologies containing the 1*_↑_*3*_↑_*2*_↑_*substructures is 15.87 among which the OFHT of topologies having the left-handed *βαβ*-unit is 1.19. (B) Provides an example of the structure containing the left-handed *βαβ*-unit. Displayed here is the 78-159 residues of chain A of the prenyl diphosphate synthase, Rv1086 (PDB ID: 2VG1). The left-handed *βαβ*-unit is colored in red. C*β* atoms of the *β*-sheet region are depicted as spheres.

In this study, We have shown that it is possible to ascertain the presence or absence of frustration in a given protein topology by considering a combination of various structural rules. Moreover, we have found that frustrated topologies exhibit low designability. This discovery has significant implications for de novo protein design, as it enables us to identify topologies with low designability based solely on their structural characteristics. For example, in a study conducted by Rocklin *et al.* [34], where more than 1,500 proteins were de novo designed with four target topologies (*ααα*, *βαββ*, *αββα*, and *ββαββ*). It was observed that the *αββα* topology presented a substantially lower success rate compared to the others. Specifically, the success rate was only 2%, while the success rates for the remaining topologies ranged from 40% to 90%. This significant difference in the success rates of designing different target topologies raises an imporatant question: what factors contribute to these disparities? We have previously reported that the *αββα* topology is a frustrated topology [22]. Given that frustrated topologies inherently exhibit lower designability, the limited success rate in designing the *αββα* topology can be attributed to its inherent low designability. Our theoretical framework not only assists in interpreting design the outcomes of design efforts but also has the potential to predict novel, designable folds [16]. As a result, it will play a crucial role in exploring the protein fold space beyond what has been sampled through natural evolution.

## 4 Materials and Methods

### 4.1 The datasets

To calculate the occurrence frequency of Homologous-groups in a topology, we utilized the ECOD database (date; 05/11/2021) [14], which contains 64,881 domains with sequence identity *<* 99%. The ECOD database classifies homologous protein domains according to categories of family and homology. The family (F) group comprises evolutionarily related protein domains with substantial sequence similarity, and the homology (H) group includes multiple F-groups with functional and structural similarities. The H-group corresponds to the superfamily in the other structural databases, such as SCOP [1] and CATH [2].

To identify the key rules for pure parallel *β*-sheet, as shown in Fig. 2, we employed the PISCES server [35] with the criteria of sequence identity *<* 25%, resolution *<* 2.5Å and R-factor *<* 1.0.

### 4.2 Analysis of *β*-sheet topologies in the dataset

In the analysis based on secondary structure assignment using the STRIDE program [15], we selectively extracted pure parallel *β*-sheets adhering to specific criteria: 1) ensuring that all adjacent pairs of *β*-strands within the *β*-sheet are parallel, 2) confirming that the *β*-sheet is open (not a barrel), 3) verifying the absence of other *β*-sheets among the *β*-strands, and 4) limiting the number of residues connecting two continuous sequential *β*-strands to fewer than 100. The process of deducing the topology from the results of secondary structure assignments followed the methodology outlined in Ref. [16].

### 4.3 Definition of the Occurrence Frequency of Homologous-group in a Topology (OFHT)

Using the ECOD dataset, we calculated the occurrence frequency of H-group in a topology *OFHT* (*T*) of a given topology *T* by summing the occupation ratio *OR*(*T, i*) of protein domains that have *T* in the *i*th H-group as

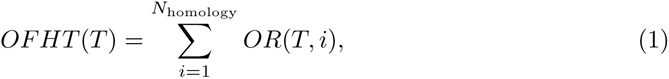

where *N*_homology_ is the total number of H-groups, and

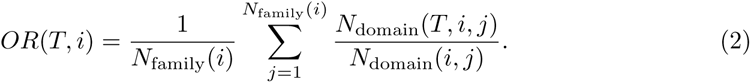

In this context, *N*_domain_(*T, i, j*) represents the number of protein domains with topology *T* in the *j*th F-group, which is part of the *i*th H-group in the dataset. *N*_domain_(*i, j*) signifies the total count of protein domains in the *j*th F-group, and *N*_family_(*i*) denotes the number of F-groups in the *i*th H-group.

## Supplemental figure captions

**S1 Fig.**
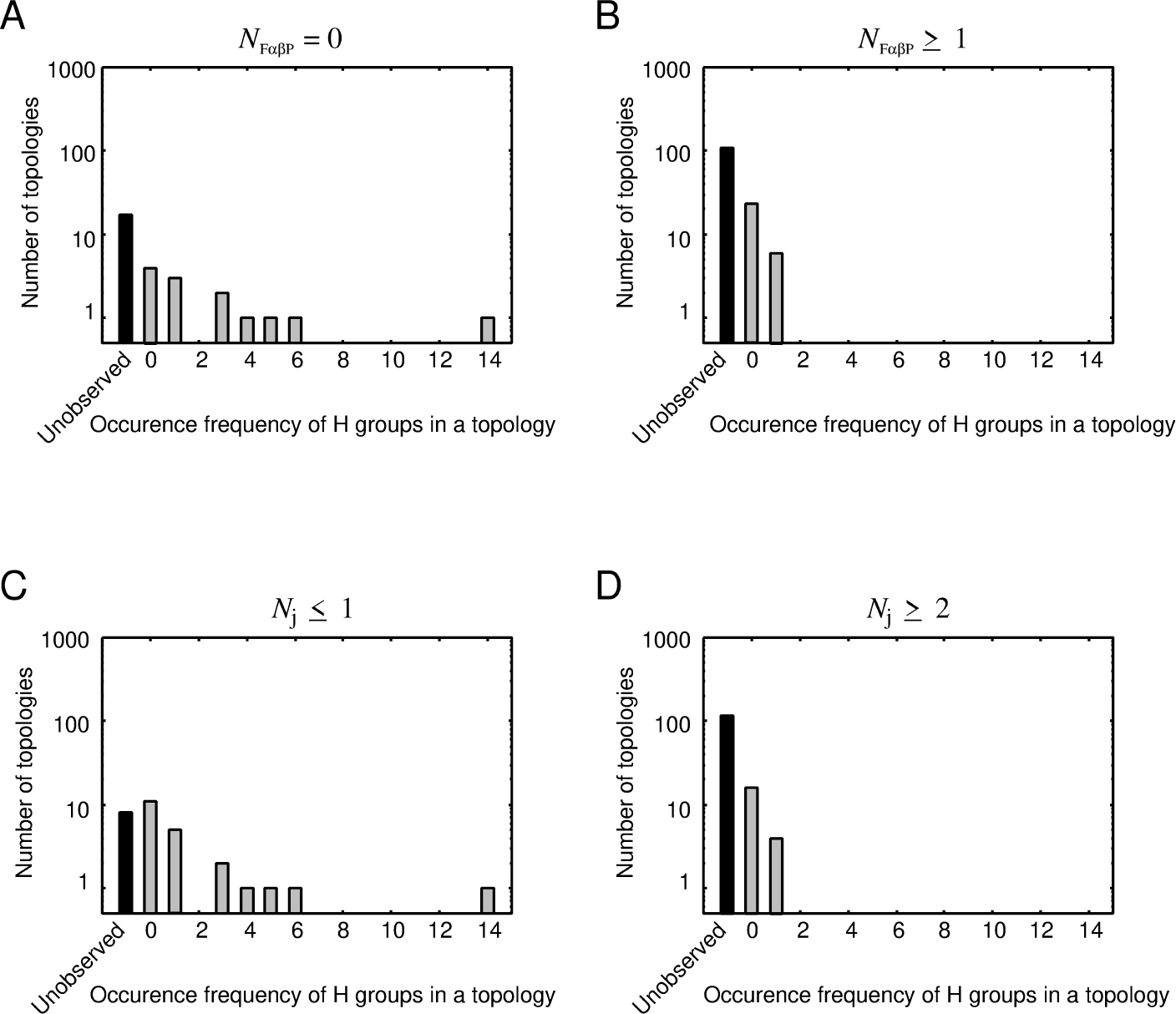
Occurring Frequency of Homologous-group in a Topology. (A) OFHT for topologies with *N*_F*αβ*P_ = 0. (B) OFHT for topologies with *N*_F*αβ*P_ *>* 0. (C) OFHT for topologies with *N_j_ ≤* 1. (D) OFHT for topologies with *N_j_ ≥* 1.

**S2 Fig.**
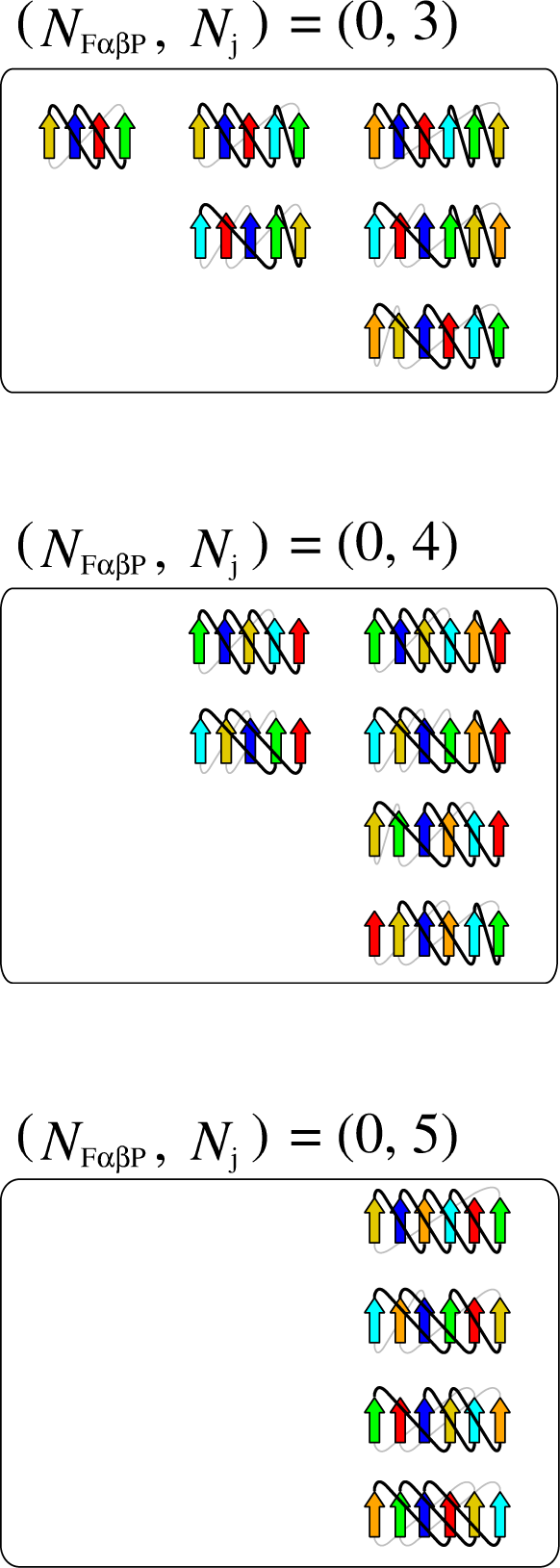
Topology Diagrams of Various States. Diagrams of the state with (*N*_F*αβ*P_*, N_j_*) = (0,3), those with (*N*_F*αβ*P_*, N_j_*) = (0,4), and those with (*N*_F*αβ*P_*, N_j_*) = (0,5).

**S3 Fig.**
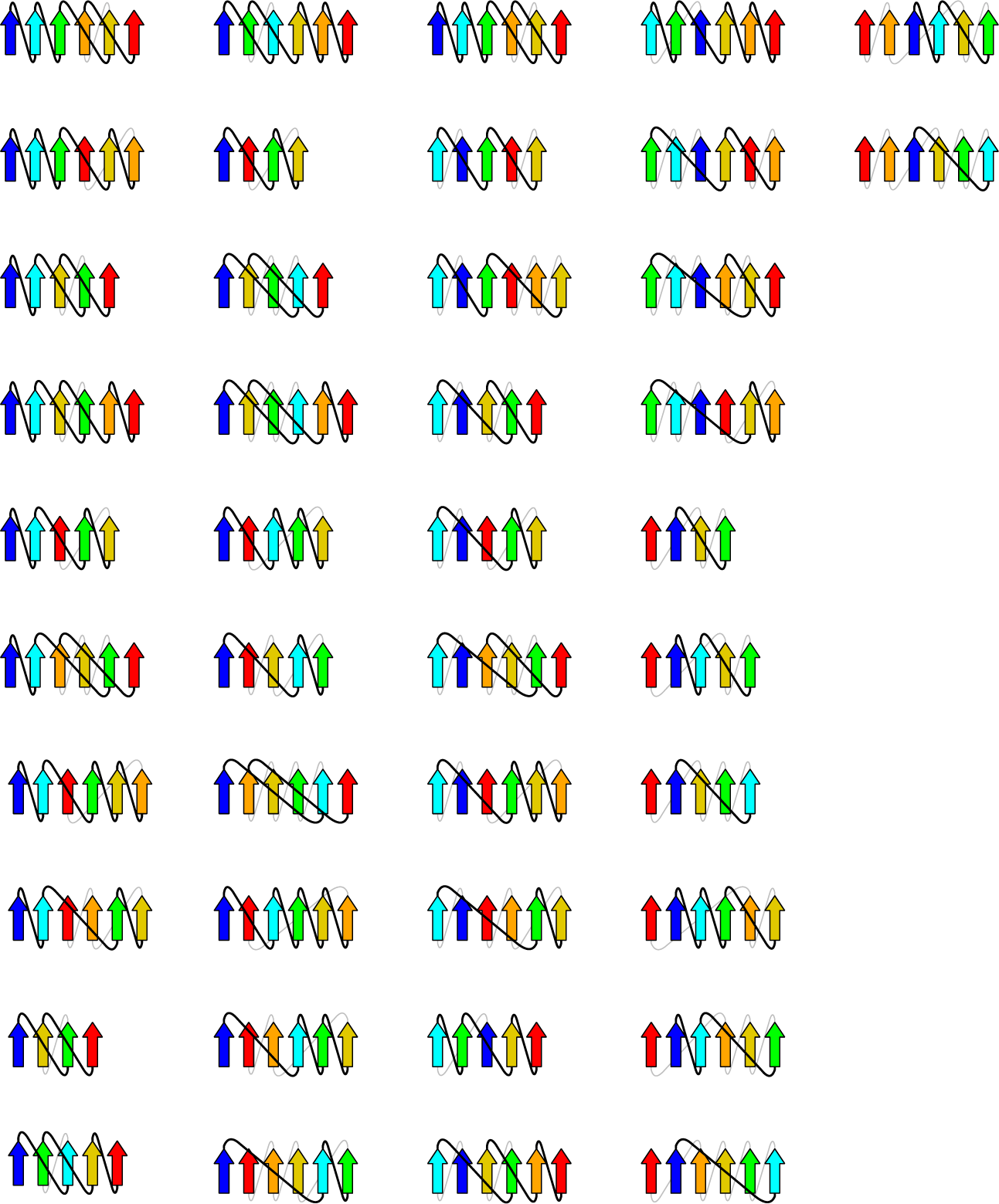
Topology Diagrams of a Specific State. Diagram of the state with (*N*_F*αβ*P_*, N_j_*) = (1,2).

**S4 Fig.**
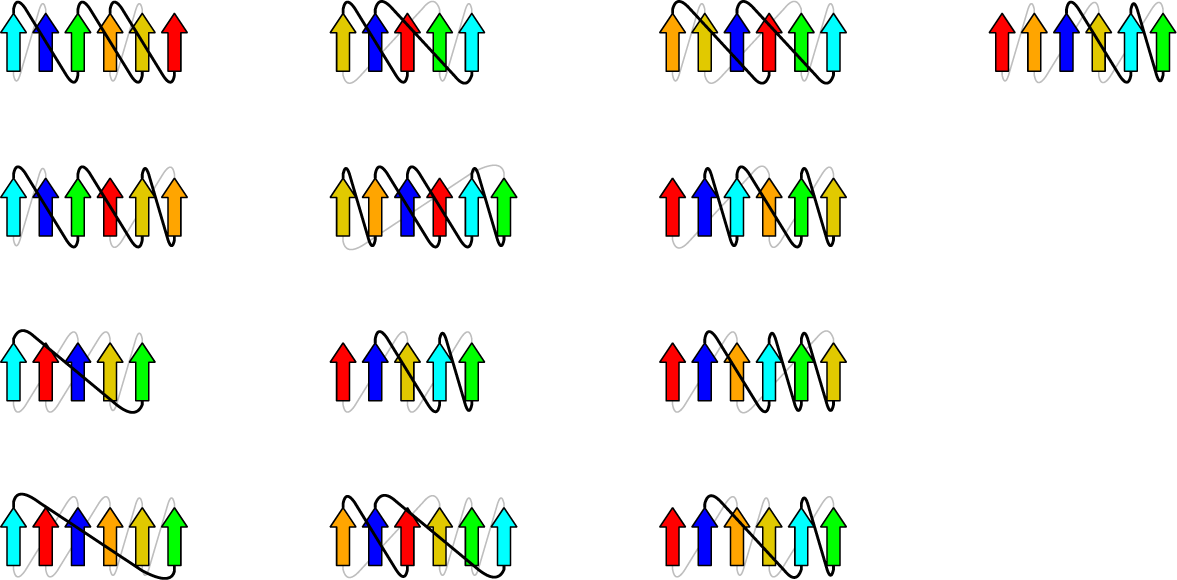
Topology Diagrams of a Specific State. Diagram of the state with (*N*_F*αβ*P_*, N_j_*) = (1,3).

**S5 Fig.**
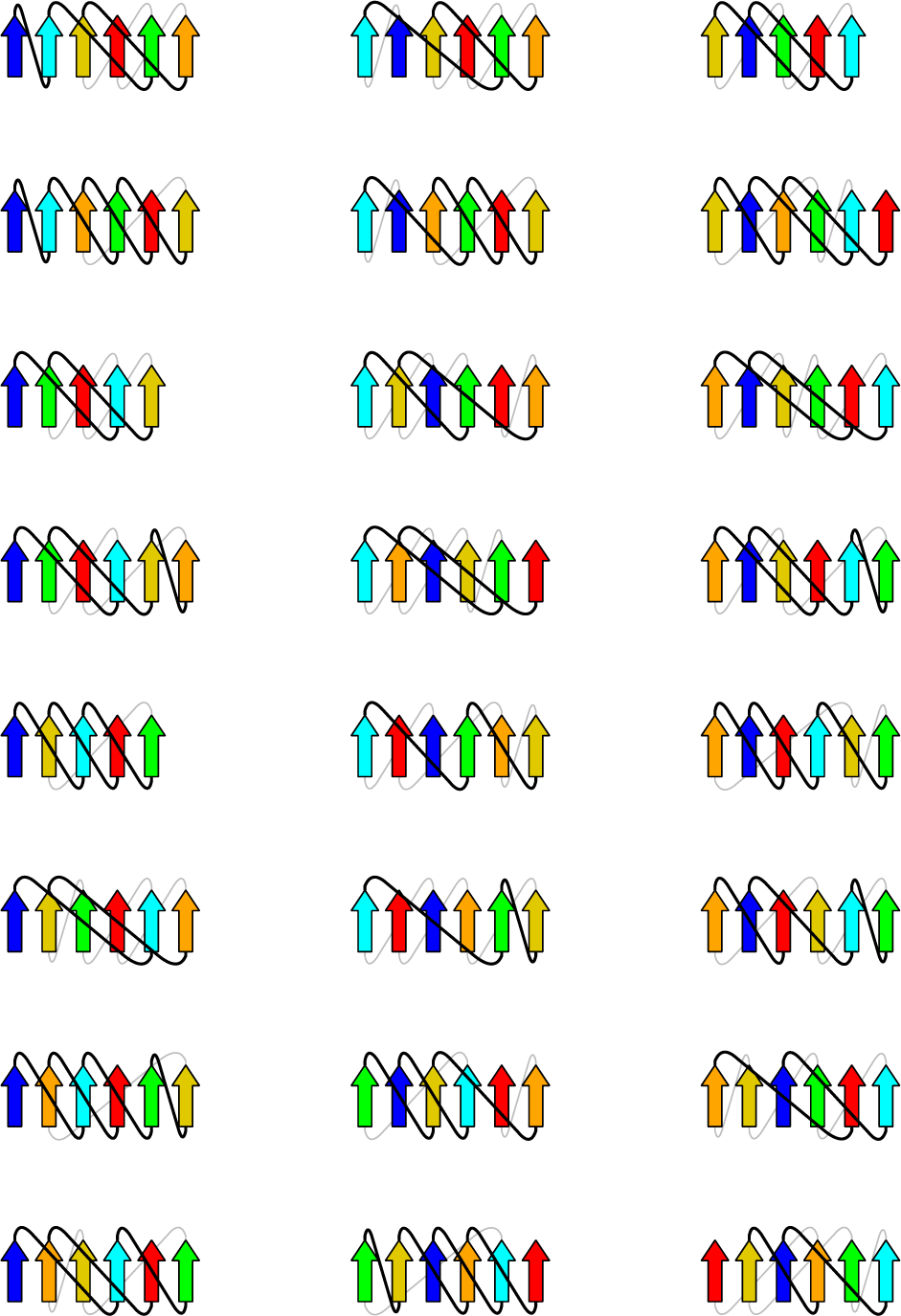
Topology Diagrams of a Specific State. Diagram of the state with (*N*_F*αβ*P_*, N_j_*) = (1,4).

**S6 Fig.**
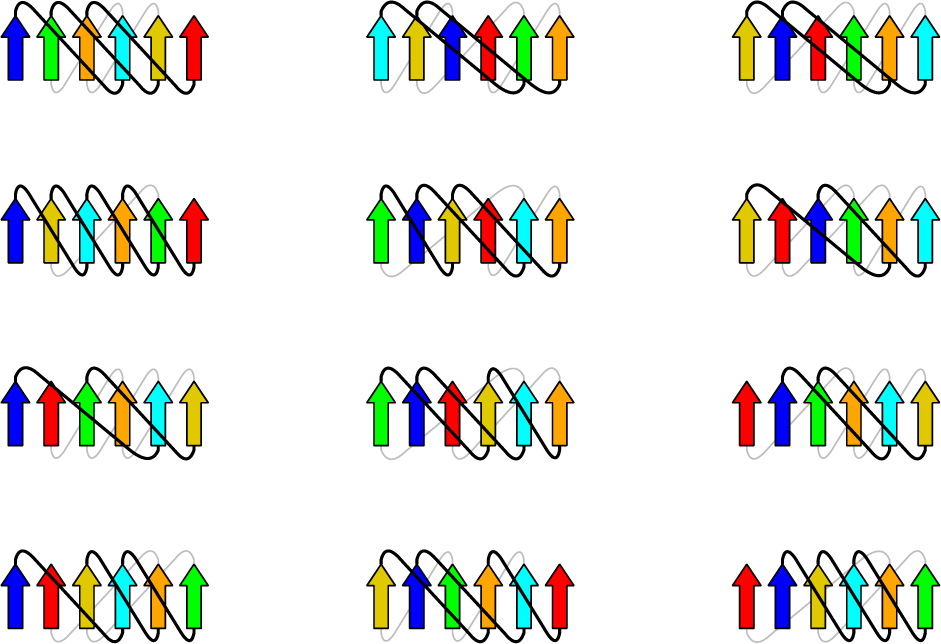
Topology Diagrams of a Specific State. Diagram of the state with (*N*_F*αβ*P_*, N_j_*) = (1,5).

**S7 Fig.**
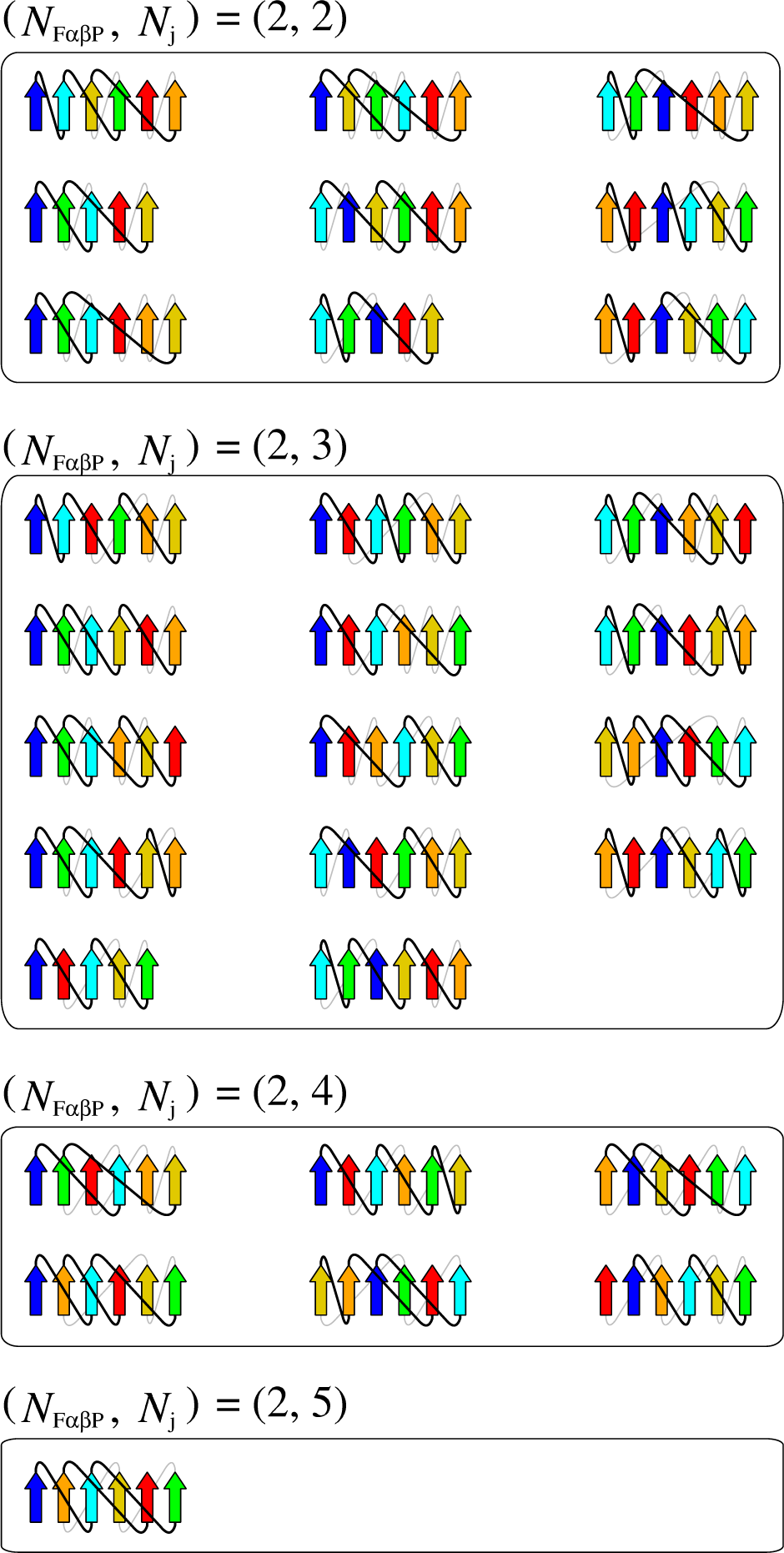
Topology Diagrams of Various States. Diagrams of the state with (*N*_F*αβ*P_*, N_j_*) = (2, 2), those with (*N*_F*αβ*P_*, N_j_*) = (2, 3), those with (*N*_F*αβ*P_*, N_j_*) = (2, 4), and those with (*N*_F*αβ*P_*, N_j_*) = (2, 5).

**S8 Fig.**
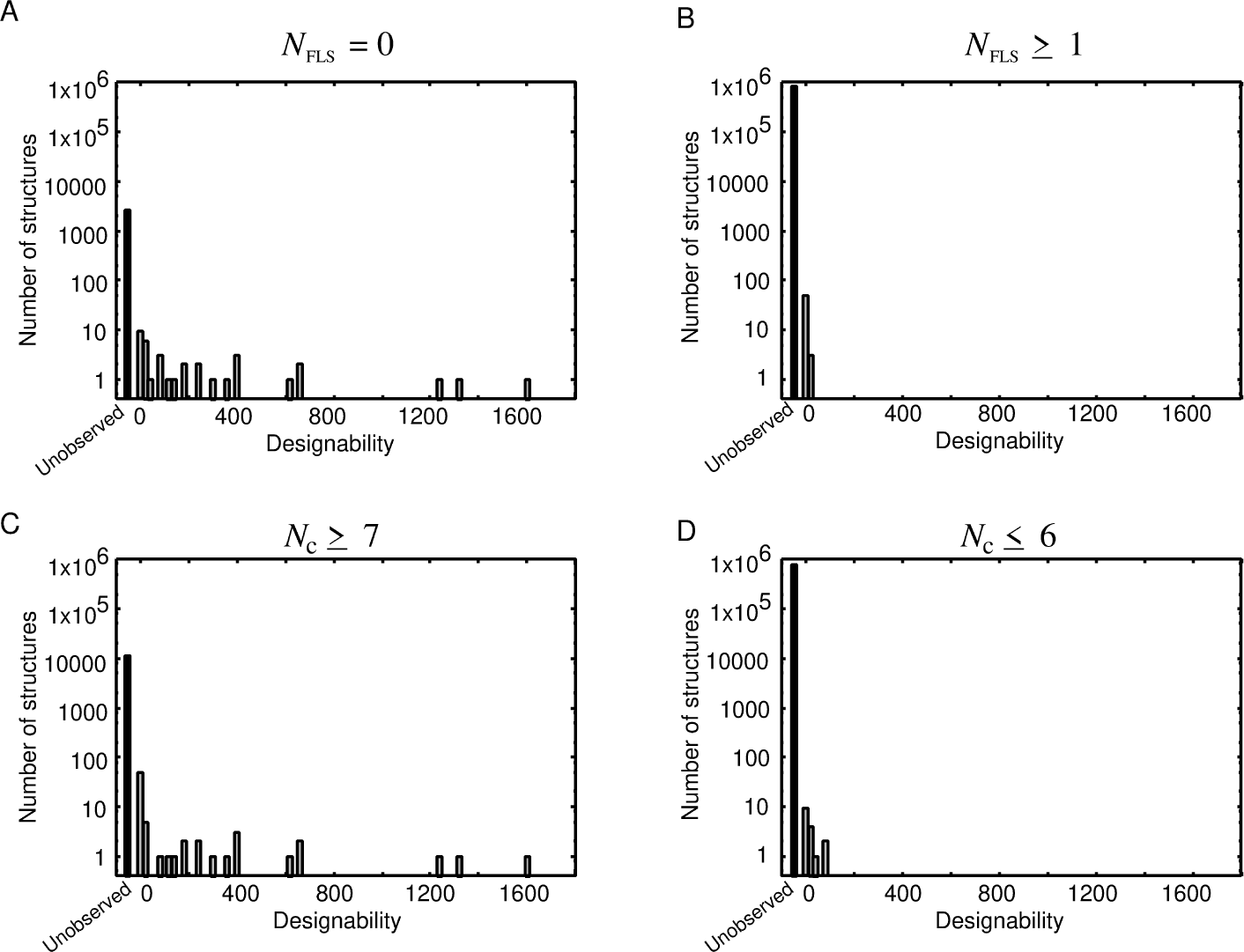
Histogram of Designability for the Lattice Model with Local Interactions. **(A) Histogram of designability for structures with** *N*_FLS_ = 0. **(B) Histogram of designability for structures with** *N*_FLS_ *>* 0. **(C) Histogram of designability for structures with** *N*_c_ *≥* 7. **(D) Histogram of designability for structures with** *N*_c_ *≤* 6.

## Acknowledgments

The authors thank Shintaro Minami and Masaki Sasai for fruitful discussions.

## Author Contributions

**Conceptualization:** George Chikenji

**Data curation:** Hiroto Murata, George Chikenji

**Investigation:** Hiroto Murata, Kazuma Toko, George Chikenji

**Methodology:** Hiroto Murata, Kazuma Toko, George Chikenji

**Project administration:** George Chikenji

**Resources:** Hiroto Murata, Kazuma Toko, George Chikenji

**Software:** Hiroto Murata, Kazuma Toko, George Chikenji

**Supervision:** George Chikenji

**Visualization:** Hiroto Murata, George Chikenji

**Writing – original draft:** Hiroto Murata, George Chikenji

